# Assessing the public health impact of tolerance-based therapies with mathematical models

**DOI:** 10.1101/073700

**Authors:** Nathanaël Hozé, Sebastian Bonhoeffer, Roland Regoes

## Abstract

Disease tolerance is a defense strategy against infections that aims at maintaining host health even at high pathogen replication or load. Tolerance mechanisms are currently intensively studied with the long-term goal of exploiting them therapeutically. Because tolerance-based treatment imposes less selective pressure on the pathogen it has been hypothesised to be “evolution-proof”. However, the primary public health goal is to reduce the incidence and mortality associated with a disease. From this perspective, tolerance-based treatment bears the risk of increasing the prevalence of the disease, which may lead to increased mortality. We assessed the promise of tolerance-based treatment strategies using mathematical models. Conventional treatment was implemented as an increased recovery rate, while tolerance-based treatment was assumed to reduce the disease-related mortality of infected hosts without affecting recovery. We investigated the endemic phase of two types of infections: acute and chronic. Additionally, we considered the effect of pathogen resistance against conventional treatment. We show that, for low coverage of tolerance-based treatment, chronic infections can cause even more deaths than without treatment. Overall, we found that conventional treatment always outperforms tolerance-based treatment, even when we allow the emergence of pathogen resistance. Our results cast doubt on the potential benefit of tolerance-based over conventional treatment. Any clinical application of tolerance-based treatment of infectious diseases has to consider the associated detrimental epidemiological feedback.

**Author summary:** Conventional therapies improve patient health by eliminating the pathogen, or, at least, reducing its burden. Recently, alternative therapies that exploit host tolerance mechanisms have received attention from the medical community as a promising strategy. These treatments aim at reducing the level of illness due to the infection, rather than eliminating the pathogen directly. Using a mathematical model, we show that although these treatments are beneficial at the individual level, they can have undesired public health consequences. In particular we show that tolerance-based treatment gives more time for the disease to spread in the population, which in turn increase its prevalence. Moreover, in the case of a low coverage of the treatment of a chronic infection, the overall mortality can increase.

## Introduction

Hosts can respond to infections in various ways. The host can reduce the pathogen replication or load and thus improve its health. In evolutionary ecology, such a response is called “host resistance”. Another possible host response is “disease tolerance”, that induces a state, in which the host, at a given pathogen load, suffers less from the negative consequences of being infected.

In evolutionary ecology, disease tolerance has received attention as a host strategy that has an impact on the evolutionary dynamics of host-pathogen systems very different to that of host resistance [1,2, 3]. In a first scenario, when resistance is cost-free, resistance genes are predicted to fix in the population [4, 5]. Another possibility, if the pathogen population is not cleared fast enough, or if there is cost of resistance, is that resistance and susceptible genes will coexist in the host population [6, 7, 8].

Tolerance genes, on the other hand, do not drive the pathogen to extinction. They even increase the prevalence of the pathogen, thus increasing the selective pressure favoring themselves. This positive evolutionary feedback often leads to the fixation of tolerance genes [9, 5] — although scenarios explaining polymorphisms in tolerance have been considered [10].

The particular molecular and immunological mechanisms that confer host resistance are clinically relevant as they provide targets for therapeutic agents. The most common treatment agents, such as antibiotics or antivirals, aim at reducing pathogen replication or burden and are often based on host resistance mechanisms. A class of agents against HIV, for example, inhibits the coreceptor CCR5 — a treatment strategy inspired by a naturally occurring polymorphism in the gene encoding CCR5 that reduces the susceptibility of individuals to HIV infection [11].

But tolerance mechanisms can also be exploited therapeutically. For example, widely used anti-inflammatory drugs reduce the negative impact of an infection without targeting the pathogen directly. Other examples of tolerance mechanisms have been found in the context of Plasmodium infections. Plasmodium infections lead to the release of heme from erythrocytes, which has proinflammatory properties. The inflammation triggers a reactive oxygen species response that can lead to liver failure in individuals with malaria. Some individuals, however, express an enzyme — heme oxygenase 1 (HO-1) — that prevents liver failure by limiting the reactive oxygen species response. Because the action of this enzyme does not affect the level of the pathogen it represents a tolerance mechanism. Inhibiting the reactive oxygen species response with a pharmacological agent that acts similarly to HO-1 has been shown to limit liver failure in mouse models [12]. HO-1 was also shown to provide host tolerance by preventing free heme from promoting severe sepsis [13]. Additional tolerance mechanisms against severe sepsis have recently been discovered [14], which involve the inhibition of cytokine production by anthracyclines. Similarly, based on the observation that sickle human hemoglobin confers tolerance to malaria, it was proposed that modulation of HO-1 via the transcription factor NF-E2-related factor 2 (Nrf2) might be a therapeutic target for treating cerebral malaria [15]. We refer to treatment strategies that are based on tolerance mechanisms as *tolerance-based treatment*.

Tolerance-based treatment is seen by some to have great promise [16] — in part, simply as a complement to more conventional treatment based on the exploitation of host resistance mechanisms. It is even hypothesized that tolerance-based treatment could hinder the development of drug resistance as it exerts a milder selection pressure on the pathogen [1, 17].

However, negative consequences of tolerance-based treatment have been considered too. Treated hosts could become healthy carriers of the pathogens, hence giving them more opportunity for transmission [9, 18, 19, 20]. Moreover, it was postulated that damage-limitation treatments — a form of tolerance-based treatment — might select for increased pathogen transmission [20, 21]. In an evolutionary context, theoretical studies also showed that pathogen virulence may increase in a tolerant host population [22]. In the context of public health, this translates into a risk of evolution towards higher virulence in response to tolerance-based treatment. But in the study by Miller et al. this higher virulence did not affect host mortality because all hosts carried the tolerance gene.

Generally, the translation of insights from evolutionary ecology to the public health situation is hindered by the fact that tolerance genes usually fix in the host population, and hosts are therefore protected from dying. Tolerance-based treatment, even if it confers great benefits to individual hosts, cannot be expected to be applied to the entire population. As a consequence, disease-induced mortality could increase when tolerance-based treatment is rolled out.

Here, we assess the promise of tolerance-based treatment (TOL) on the population level and investigate the public health impact of treatment at various levels of coverage. While most previous studies focused on changes in the incidence and prevalence of the disease, we specifically focus on the disease-induced mortality. To this end, we use mathematical models. Specifically, we extend the well-known epidemiological SIR model, featuring susceptible, infected, recovered individuals, by an additional compartment of treated individuals. We investigate the effect of TOL for two types of infections: acute and chronic.

While, at the individual level, TOL can be effective, and may even prevent the evolution of pathogen resistance in the long run, it is problematic epidemiologically. Indeed, we show that for chronic infections, the disease-induced mortality increases for low treatment coverage, because of the introduction in the population of asymptomatic carriers of the disease. Comparing TOL to conventional treatment, based on a reduction of load (ROL), we find that for both, acute and chronic infections, ROL always outperforms TOL even when we consider that pathogen resistance can emerge against ROL.

## Methods

### Compartment model for a single treatment

We consider an extended SIR model in which infected patients can be treated. The model is depicted in Fig 1. Susceptible (uninfected) hosts *S* enter the population at a rate Λ, and die at a per capita rate *δ*. They can be infected by individuals that are either untreated (*I*) or treated (*T*). Susceptible hosts become infected at a rate that depends on the number of susceptible *S*, the total number of infected *I* +*T* and the transmission coefficient *β*. This reflects contact-dependent transmission from infected hosts to uninfected hosts. Infected hosts die at a rate *δ* + *v*, with *v* ≥ 0 indicating a higher mortality of infected than susceptible. Infected can recover at a per capita rate r and become part of the recovering population *R*. Similarly, treated infected die at a rate *δ* + *v*_*T*_ and can recover at a per capita rate *r*_*T*_. Untreated infected individuals are treated at a rate *θ*, and become immediately treated infected.

**Figure 1:**
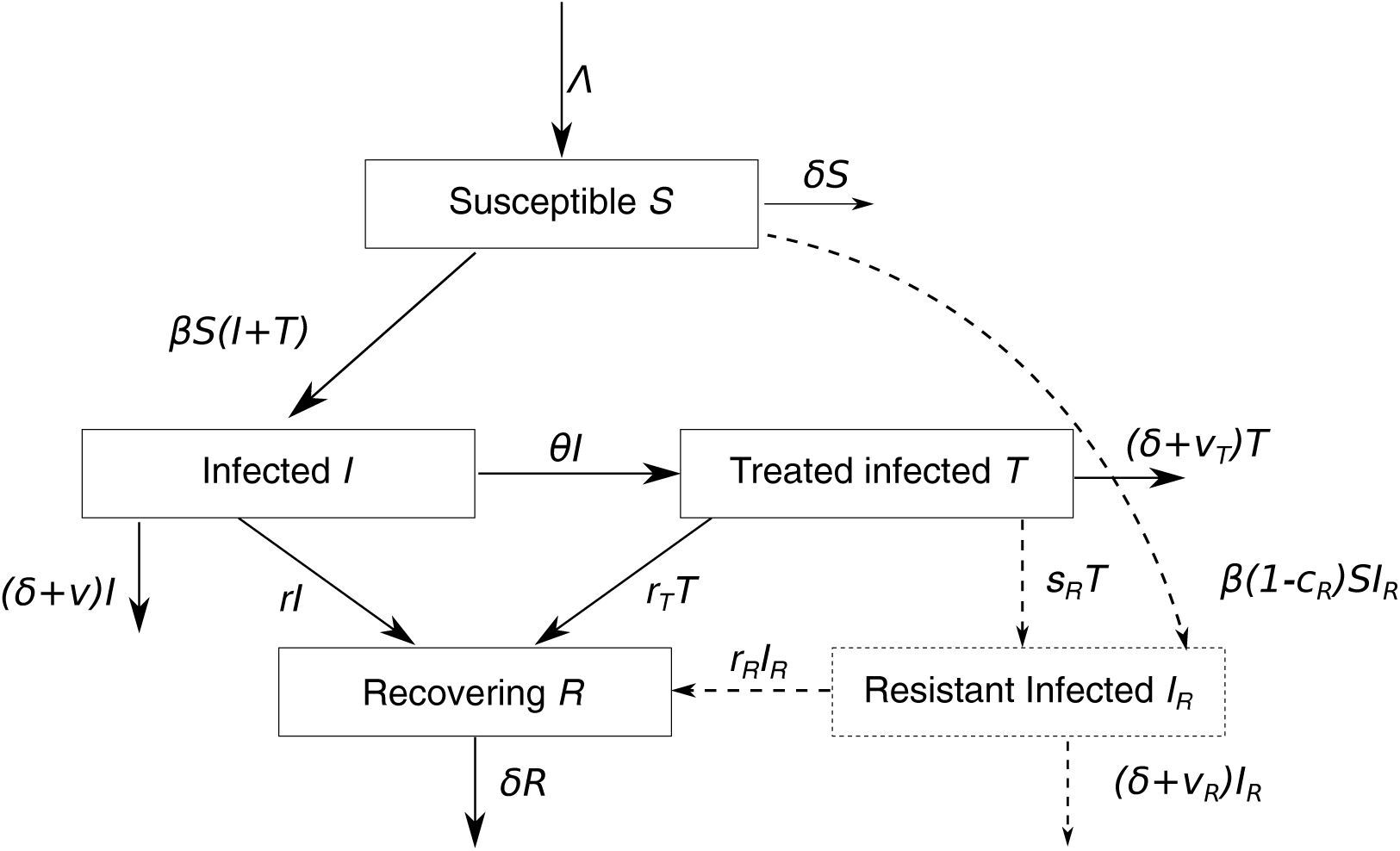
Schematic of the compartment model. The main model is presented in eq. (1). The extended model, that includes resistance to treatment, is represented in dashed lines in the schematic and is detailed in eq. (6).

The model is summarized by the system of ordinary differential equations

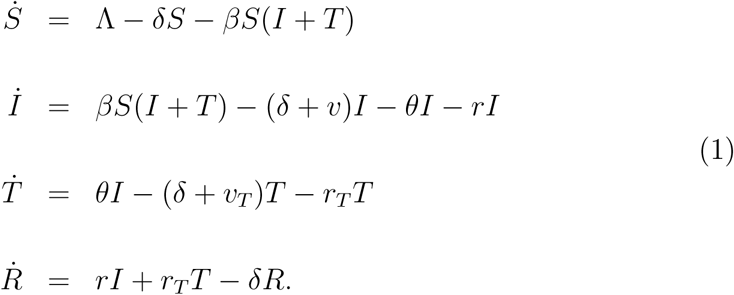

We assume that treatment is instantaneous and that infected individuals become treated infected at a rate *θ*. Infected individuals leave the infected state at a rate *δ* + *v* + *r* + *θ*, and a fraction

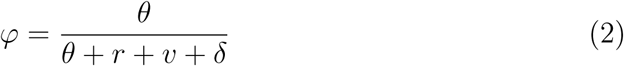

of them become treated infected. This rate ratio can be regarded as the probability for an infected individual to be treated instead of leaving to another state (death or recovery).

### Implementation of the treatments

For the implementation of TOL, we assume that the death rate is lower for treated than untreated individual (*v*_*TOL*_ = *v*_*T*_ < *v*), but that the hosts are still infectious (*r*_*TOL*_ =*r*_*T*_ = *r*).

For a ROL the hosts recover faster than without a treatment, but the mortality rate is the same (*r*_*ROL*_ =*r*_*T*_ > *r* and *v*_*T*_ = *v*).

In addition, we consider a treatment that combines the properties of TOL and ROL (TOL+ROL). We translate this effect by combining the benefits of the two treatments, the increased recovery rate of ROL and the decreased mortality rate of TOL (*r*_*TOL*+*ROL*_ = *r*_*ROL*_ and *v*_*TOL*+*ROL*_ = *v*).

In this study, we will evaluate the benefits of a treatment by considering three measures. First, the *incidence*, defined as the number of new infections in a period of time, is

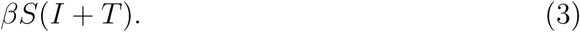

Second, the *prevalence* of the disease, which is the ratio of infected individuals over the total population, is

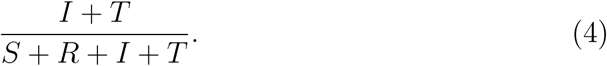

Finally, we will measure the *disease-induced mortality*, which is the fraction of deaths that are due to the disease over the total number of deaths, given by

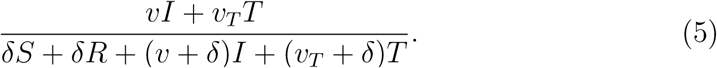

Throughout this study, we evaluate how these three measures vary at equilibrium with the fraction of treated population. Model (1) has two equilibria: the disease free equilibrium (DFE) and the endemic equilibrium (EE). Whether one equilibrium or the other is attained depends on the relative value of the model parameters (See Supplementary Text S1 for details on the equilibrium values and the equilibrium stability analysis).

Moreover, the efficacy of the treatments are evaluated for two types of infection: an acute infection, which is highly transmissible, with fast death and recovery rates, and a chronic infection, with a slower dynamics than the acute infection, and for which no recovery is possible in the absence of treatment.

### Model of pathogen resistance to the conventional treatment

In an extension of the model, we assumed that ROL can induce pathogen resistance to treatment, and added to the compartment model the number *I*_*R*_ of hosts that are infected by pathogens resistant to treatment. Pathogens infecting treated individuals can develop resistance to treatment at a rate *s*_*R*_, called acquired (or de novo) resistance. Susceptible individuals can be infected by treated, untreated, or resistant hosts. If a susceptible individual is infected by a non-resistant, it becomes infected non-treated, but becomes resistant if it is infected by a resistant individual. Individuals infected by the resistant pathogen die at a rate *δ* + *v*_*R*_. The model is an extension of model (1) with an additional compartment for the infected resistant and the corresponding transition rates (Fig 1) and is described by the set of equations

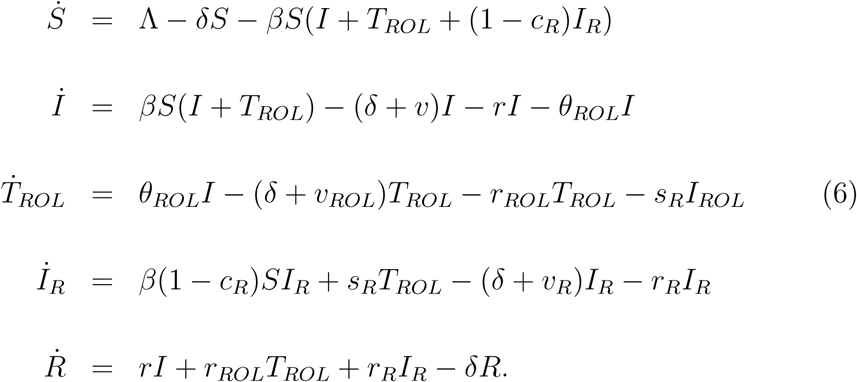

In our model, resistance can appear in two ways. First, individuals receiving ROL can acquire resistance at a rate *s*_*R*_, and second, susceptible individuals can be infected by resistant infected. Transmission of the resistant pathogen is described by a mass action law with a transmission coefficient *β*(1 — *c*_*R*_), where *c*_*R*_ > 0 represents the fitness cost of transmission associated to pathogen resistance to treatment.

Similarly to the case without resistance to treatment, the system of equations has a disease-free and an endemic equilibrium. Additionally, there is a third equilibrium where all the infected individuals are resistant to the treatment. It can be shown analytically that there is always a threshold value of the treatment coverage such that only resistant pathogens exist at equilibrium, provided that the cost of transmission of the resistant pathogen *c*_*R*_ is small enough. Details of the equilibrium values for the model of pathogen resistance are given in Supplementary Text S2.

### Choice of parameters

#### Parameters in the absence of treatment

The death rate without a disease is set to *δ* = 1/70 years^−1^ and *Λ* is determined so that the ratio Λ/*δ* is equal to a fixed population size in the case of no epidemic. We choose *Λ* = 10^6^*δ* = 1.43 10^4^ years^−1^. The transmission coefficient *β* is chosen so that the basic reproductive ratio 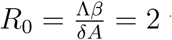 when no treatment is applied. We assume that TOL reduces the death rate of the disease, and that ROL increases the recovery rate.

#### For an acute infection

The parameters *r* and *v* are chosen so that one out of fifty infected individuals dies without treatment. Hence, 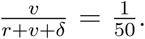

Moreover, we set the duration of the infection to be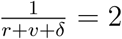

- Death rates *v* = 0.01 week^−1^ *v*_*ROL*_ = *v*_*R*_ = *v, v*_*TOL*_ = 0.
- Recovery rates *r* = 0.49 week^−1^, *r*_*ROL*_ = 0.98 week^−1^, *r*_*TOL*_ = *r*, *r*_*R*_ = *r*.
- Transmission coefficients *β* = 1.0 10^−6^ week^−1^, fitness cost of pathogen resistance to treatment *c*_*R*_ =0.2.
- Rate of acquired resistance *s*_*R*_ = 0.007 week^−1^.

#### For a chronic infection

Infected untreated individuals die from the infection in 10 years. They do not recover from the infection in the absence of treatment, and in one year with ROL.

- Death rates *v* = 0.002 week^−1^, *v*_*ROL*_ = *v*_*R*_ = *v*, *v*_*TOL*_ = 0.
- Recovery rates *r* = 0, *r*_*ROL*_ = 0.02 week^−1^, *r*_*TOL*_ = 0, *r*_*R*_ = 0.
- Transmission coefficients *β* = 4.4 10^−9^ week^−1^, fitness cost of pathogen resistance to treatment *c*_*R*_ = 0.2
- Rate of acquired resistance *s*_*R*_ = 0.007 week^−1^.

In the Supplementary Information, we provide results for various sets of parameters (Figs S1, S2). We consider an extension of the model with a cost of treatment associated to treatment (Fig S1). We also consider various values of *R*_0_ (Fig S2). In both cases, the main conclusions of this study remain unchanged. In another extension, we treat the case of imperfect treatment, characterized by 0 ≤ *v*_*TOL*_ ≤ *v* (Supplementary Text S3 and Fig S3).

## Results

We investigated the impact of TOL on the population of hosts with a mathematical model. We based our model on the SIR model [23, 24, 25], which describes susceptible, infected, and recovered hosts. To describe treatment, we divided the compartment of infected individuals into a treated and an untreated compartment (see Fig 1). Treated individuals arise from infected untreated individuals at a given rate and remain infectious.

ROL treatment is implemented in this model as an increased recovery rate of treated individuals. Treated individuals are less infectious than untreated ones because they recover faster. Per unit time their infectiousness is not assumed to be affected by treatment. Tolerance-based treatment, on the other hand, does not affect the rate of recovery but lowers the disease-induced death rate of treated hosts. Because TOL, by definition, does not affect the pathogen load we assume that treated individuals are equally infectious as untreated ones per unit time. However, they cause more infections than untreated individuals because they live longer and thus have an extended infectious period.

We assess the effect of treatment on three epidemiological quantities: prevalence, incidence, and disease-induced mortality. We determine these quantities in the endemic equilibrium for different levels of treatment coverage. In the Method section, we show how we calculated these quantities from our model equations. Because TOL increases the infectious period, we expect the incidence and prevalence to rise. The effect of TOL on the population-wide disease-induced mortality depends on how the higher prevalence is balanced by the lower mortality of treated individuals.

We investigate the impact of TOL and ROL for acute and chronic infections. Acute infections are modeled with influenza in mind, and are characterized by a short period of infection and a high transmission. Specifically, an untreated infection is assumed to last two weeks and the case fatality proportion is set to 1/50. Chronic infections are assumed to last years and there is no recovery, as for HIV infection. For both types of infection, the basic reproduction number *R*_0_ = 2 in the absence of treatment. We neglect seasonal fluctuations in any of the model parameters because this would preclude an equilibrium analysis.

### Tolerance-based treatment is beneficial on the population level only when coverage is large

We assessed the efficacy of treatment by evaluating numerically the incidence, prevalence and disease-induced mortality for different levels of treatment coverage. We vary treatment coverage by changing the rate of treatment, *θ* We define treatment coverage as the fraction of treated hosts, which is the product of the rate of treatment times the duration of an untreated infection:
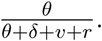

ROL reduces the incidence, the prevalence and the disease induced mortality (Fig 2a-f). Thus, ROL is unambiguously effective on the population level, which is owed to the fact that it shortens the infectious period. For the parameters chosen here, chronic infections can even be eradicated by ROL if the coverage exceeds 55% (Fig 2d-f).

**Figure 2:**
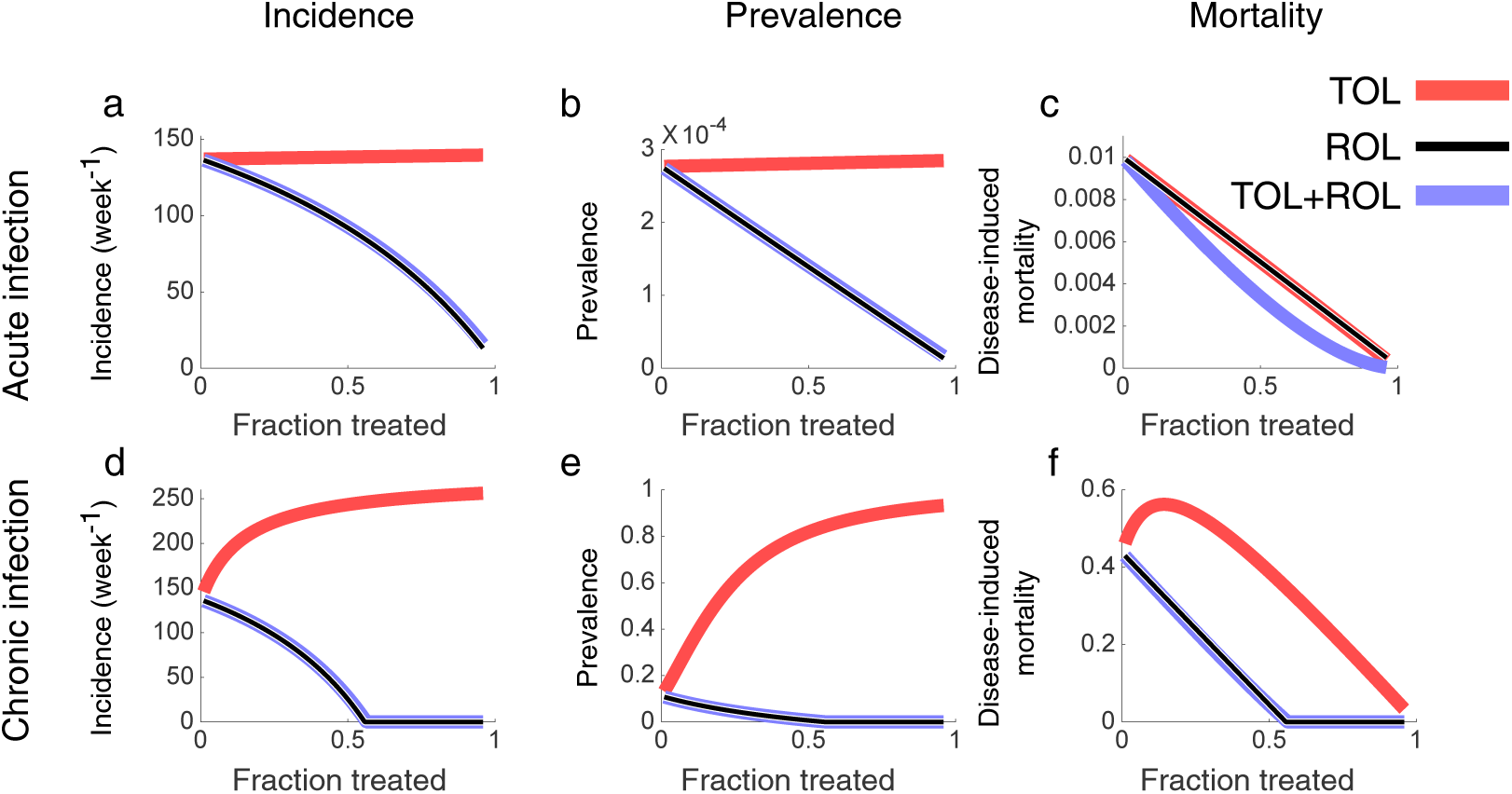
Evaluation of the efficacy of TOL and ROL in acute and chronic infections. Equilibrium values of the incidence (a,d), prevalence (b,e), and fraction of mortality due to infection (c,f), which normalizes the disease-induced mortality by the overall mortality. These epidemiological measures are plotted against the fraction of the population that receives treatment. The effect of TOL is displayed in light red, that of ROL in black, and that of TOL and ROL combined in blue. Parameters for both acute and chronic infections are given in the Methods section.

TOL, on the other hand, affects the epidemiology very differently. Unlike ROL, TOL increases incidence and prevalence (Fig 2a,b,d,e). This effect is due to TOL lengthening the infectious period. The rise is much less pronounced for acute than for chronic infections. For the acute infection, the incidence increases by only 2% with the treatment coverage (Fig 2a), while, for chronic infections, it rises from 140 to 260 new infections per week (Fig 2d). Similarly, the prevalence increases by 2% in acute infections (Fig 2b), as compared to the chronic infection where the prevalence rises from 0.11 to 0.93 (Fig 2e). The more pronounced effects of TOL for chronic versus acute infections are due to recovery rate being much lower than the death rate in chronic infection. An analytical explanation is provided in Supplementary Text S4.

In acute infections, the effect of TOL on disease-induced mortality is similar to ROL. For large treatment coverage, disease-induced mortality is reduced with both treatment approaches. In chronic infections, however, TOL can even increase the disease-induced mortality when the coverage is low (Fig 2f). This is due to the fact that there is no recovery from chronic infections in our model, and treated hosts keep infecting for life. This effect is maintained even when infected hosts can recover, provided that the time to recovery is long enough. Thus, for chronic infections, the population-level disadvantages of TOL outweigh the benefits to the individual.

### Combining ROL and TOL decreases the mortality in acute infections

To assess if TOL could be a useful addition of our treatment repertoire when combined with ROL, we implemented a strategy we call TOL+ROL. Individuals receiving this combination treatment experience both, faster recovery (due to ROL) and decreased mortality (due to TOL) (see the system of equations (1) in the Methods section). Again, we calculate the endemic incidence, prevalence and disease induced mortality under TOL+ROL and compare it to these measures under TOL and ROL alone (Fig 2).

We find that TOL+ROL does not improve on ROL in terms of incidence and prevalence but it does not do worse either — a very conceivable outcome given that individuals receiving TOL+ROL live longer. Apparently, the gain in life expectancy of individuals receiving TOL+ROL does not translate into a substantial increase of incidence and prevalence (Fig 2a,b,d,e) because, due to fast recovery under TOL+ROL, this strategy does not produce many asymptomatically infected individuals that increase the force of infection. With respect to disease induced mortality, TOL+ROL outperforms ROL in acute infections (Fig 2c). This effect is a direct consequence of the lower mortality rate of individuals treated with TOL+ROL as compared to ROL.

Thus, TOL can be a useful additional treatment strategy, especially for acute infections, if combined with ROL. Unlike on its own, in combination with ROL it is certainly not predicted to have negative public health consequences. It can be shown that TOL+ROL has public health benefit across the entire range of possible treatment coverage if the faster recovery outweighs the increase in life expectancy. Formally, the duration of infection in treated individuals, 1/(*δ* + *v*_*T*_ + *r*_*T*_), needs to be smaller than that in untreated individuals, 1/(*δ* + *v* + *r*).

### Resistance to ROL treatment

Up to this point in our analysis, TOL did not have any advantage over ROL. One supposed advantage of TOL, which we have not yet taken into account, is that it does not provoke pathogen resistance. The reason for this is that TOL does prolong rather than shorten the infectious period in treated individuals and thus increases pathogen fitness. Reduction of the pathogen load that follows ROL treatment, on the other hand, imposes a negative selection pressure on the pathogen by reducing the duration of infection. In response, the pathogen might evolve resistance.

To assess the promise of TOL more comprehensively, we included pathogen resistance to ROL into our model. To this end, we added a compartment for individuals infected with resistant pathogen strains (Fig 1). Individuals enter this compartment either after being infected and receiving treatment, which may trigger de novo resistance emergence. Resistant pathogen strains are also assumed to be transmitted. We further assume that resistant pathogens render ROL ineffective, i.e. individuals infected with resistant pathogen strains have the same recovery and disease-induced death rates as untreated infected individuals. Lastly, we allow resistant pathogens to carry a fitness cost in terms of a lower transmission coefficient.

We do not implement the emergence of resistance as a stochastic process. While this would be admittedly more realistic, treating resistance deterministically is more favorable to TOL. We thus present the best case scenario for TOL.

We find that resistance outcompetes the wildtype pathogen strain if the treatment coverage exceeds a threshold, *φ*_*ROL*_. For the parameters we chose, *φ*_*ROL*_ = 0.2 for acute and *φ*_*ROL*_ = 0.4 for chronic infections (Fig 3a and e). Below this threshold, wildtype and resistant pathogen strains coexist.

**Figure 3:**
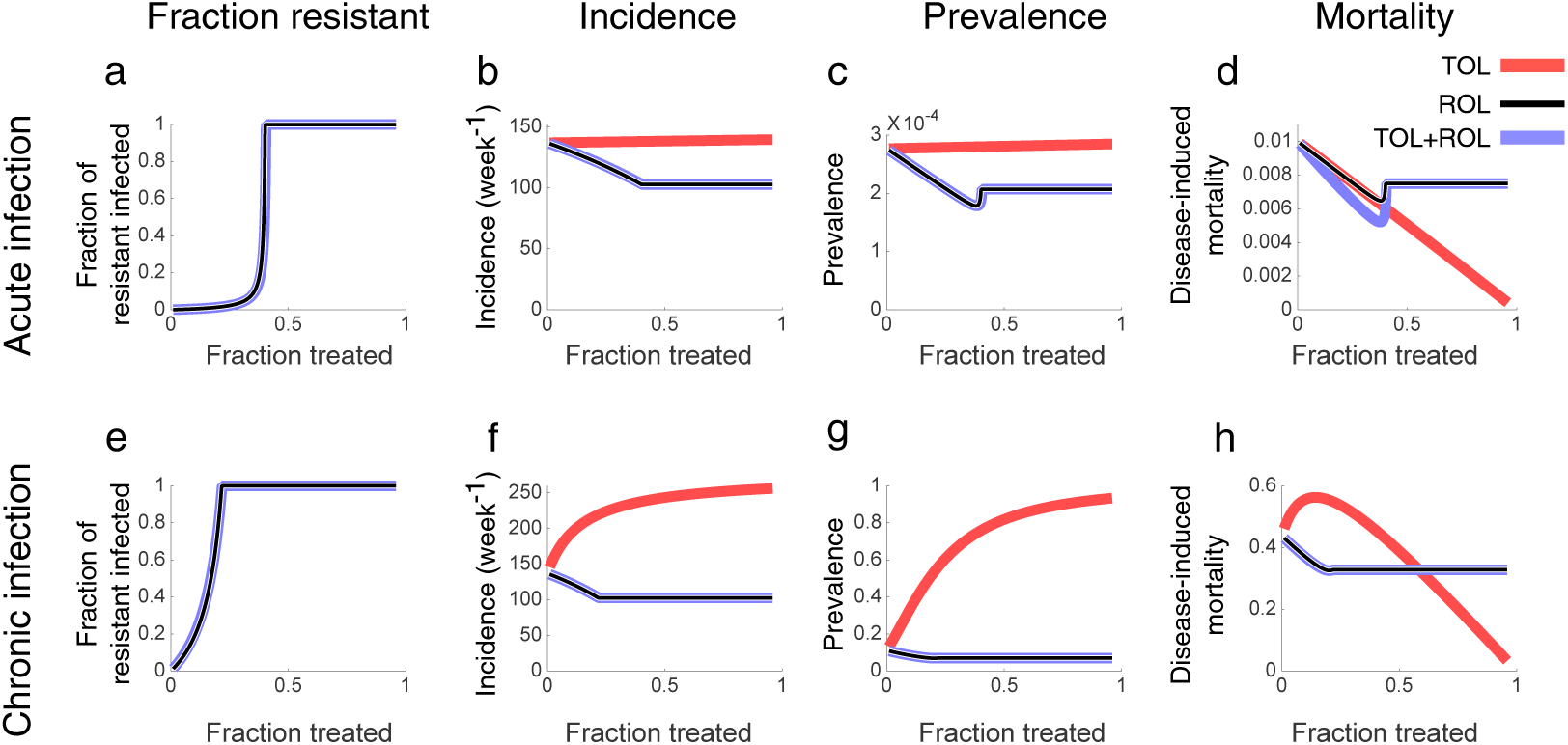
All pathogens become resistant to treatment for large coverage of the ROL treatment. Fraction of the population infected by the resistant pathogen over the total infected population (a, e), and equilibrium values of the incidence (b,f), prevalence (c,g), fraction of mortality due to infection (d,h). These epidemiological measures are plotted against the fraction of the population that receives treatment. The effect of ROL is displayed in black, that of TOL and ROL in blue. TOL, which does not lead to resistance, is represented in red and is reproduced for comparison from Fig 2. Parameters for both acute and chronic infections are given in the Methods section.

Below the threshold *φ*_*ROL*_, the incidence, prevalence and disease-induced mortality are very similar to the model without pathogen resistance. Above the threshold, the three measures under ROL and TOL+ROL become independent of treatment coverage because treatment is assumed not to affect resistant pathogen strains. The incidence and prevalence under ROL and TOL+ROL still remain below the levels they attain under TOL even if we allow pathogen resistance to evolve. However, the disease-induced mortality under TOL can become smaller than under the other treatment strategies for high treatment coverage (Fig 3d and h). For chronic infections, the levels of treatment coverage for which TOL becomes advantageous is much larger than the threshold *φ*_*ROL*_: TOL becomes beneficial if treatment coverage exceeds approximately 60% (Fig 3h). Some of these results depend critically on our equilibrium assumption (see Discussion).

## Discussion

Tolerance mechanisms are currently considered to be exploited therapeutically [16]. Undoubtedly, once developed, tested and approved, TOL will benefit treated individuals. This clear benefit of TOL should not cloud our view for what is most important: the direct public health consequences of such treatment as measured by the number of lives saved population-wide. In this paper, we assessed the potential of TOL from the public health perspective.

We find that TOL is beneficial at the population level for acute infections. However for chronic infections, irrespective of whether we take pathogen resistance into consideration, TOL on its own is not a promising treatment strategy from a public health perspective. The levels of coverage required to make mortality lower for TOL than in the absence of treatment — above 50% in our simulations — could probably be attained only for chronic infections in countries with an excellent public health infra-structure. However, if treatment is not perfect (Supplementary Text S3 and Fig S3), consequences of TOL are even more dramatic. The epidemiological feedback leads to an higher mortality with treatment than in the absence of treatment.

TOL is also of limited use as a supplement to other interventions. We considered the combination of TOL and ROL. Except for acute infections and low coverage, this combination is not better than ROL on its own.

Because specific agents for tolerance-based treatment are still in development, a mathematical modeling approach is currently the most appropriate way to assess the public health impact of TOL. Mathematical modeling also enables us to gain insights into a wide range of different infections and to apply TOL and ROL at different levels of coverage, separately and in combination. Having said that, experimental study systems are being developed that will allow the direct assessment of the epidemiological effect of TOL. A recently developed transmission model, involving Salmonella infection of tolerant and non-tolerant mice, highlighted the role of tolerant mice in the spread of the infection [26]. While being certainly more biological, such transmission models cannot easily be scaled up to the population sizes a pathogen encounters during an epidemic. Furthermore, the treatment agents might have multiple effects that do not easily fall into the categories of TOL or ROL. Thus, even with transmission models, mathematical modeling will play a role in assessing the public health consequences of treatment.

There is a rich literature on the evolutionary ecology of disease tolerance (as reviewed by Boots et al 2009). Most mathematical models in this context focus on the evolution of tolerance as a host trait. In these models, tolerant hosts are competed against resistant hosts under pathogen pressure. The fact that tolerant hosts lead to an increase of pathogen prevalence leads to a benefit of tolerant over resistant or non-tolerant hosts. Thus, unlike in our model of tolerance-based treatment, the epidemiological feedback “biases competition” [5] in favor of tolerance as a host strategy.

A few of the evolutionary models consider the evolution of pathogens in response to host resistance or tolerance [5, 20, 22, 27, 28]. Miller et al, 2006 for example, investigate pathogen evolution in host populations that exhibit a variety of tolerance mechanisms. They find that the “absolute mortality”, i.e. the total number of deaths per unit time, can increase for virulent pathogens with the level of tolerance for both, unevolved and evolved pathogens. But the “relative mortality” — that is adjusted for the host population size and is comparable to the mortality we consider — is not found to increase (see their Fig 6). It is important to note, that in their study all hosts are tolerant, which in our model would correspond to 100% treatment coverage. The same applies to the study by Vale et al, 2014 that focused on the evolutionary response of the pathogens to various damage-limitation treatments and found that prevalence can increase and the pathogen can evolve higher virulence. While identifying important negative evolutionary consequences of TOL, these studies did not identify the conditions under which the application of tolerance-based treatment would lead to an epidemiological increase in mortality under realistic levels of treatment coverage. Of note, some models [29] consider the coevolution of tolerance in hosts and pathogen virulence. In this case, they obtain that in some cases that tolerance is not a trait shared by the whole population, but is variable in the host population.

Formally, our modeling of TOL bears most similarities with studies on the epidemiological and evolutionary aspects of imperfect vaccines [30, 31]. In particular, the anti-toxin vaccines discussed in these studies reduce the virulence in the vaccinated hosts without affecting transmission, recovery, or infection probability, and are therefore identical to TOL in terms of their effect on the infection parameters of an individual. However, anti-toxin vaccines differ from TOL in that they are given also to uninfected hosts. Thus, these vaccines are equivalent to prophylactic TOL. The major difference between prophylactic tolerance-based treatment and TOL is the number of people that need to be treated to reach similar effects. In particular, to reach a substantial decrease in the overall mortality in the context of chronic infections, the prophylaxis of a large fraction of the entire population is required. This is described in more details in Supplementary Text S5, where we establish a model of prophylactic TOL.

In contrast, our analysis focuses on a treatment given only to infected individuals, and is thus not prophylactic. Moreover, the treatment is given to a fraction of the infected population. Thus, our model addresses the impact of treatment coverage on the epidemiological feedback of TOL. In particular, we focused on the disease-induced mortality — the most relevant entity for public health for severe infections. This feedback is most pronounced at low treatment coverage where the increase in prevalence due to TOL meets a sufficient frequency of untreated, and hence vulnerable, hosts.

TOL could be useful when linked with transmission control, or if the tolerance induction goes hand-in-hand with a reduction of transmission. Sometimes such transmission reductions are a side effect of TOL [32]. Our model allows us to calculate the transmission reduction required for TOL not to increase disease prevalence: transmission has to be reduced by 2% in acute and by 88% in chronic infection (see Supplementary Text S6). The substantial reduction required for chronic infections is difficult to achieve and, once again, TOL, even with transmission control, might be useful only against acute infections (Supplementary Text S7 and Fig S4). However, such transmission control might reduce pathogen fitness, and pathogen could evolve in response to TOL. While we did not consider the potential evolutionary consequences of TOL in our study, recent papers [20, 21] suggest that TOL could increase pathogen virulence, especially when virulence and pathogen fitness are tightly linked [33].

But these considerations about transmission control might still be too optimistic. It has been shown that antipyretic treatment - that aim at reducing fever during flu infection - can increase transmission [34]. This is due in part to more frequent contacts between an individual with reduced fever and susceptibles, and also to the lengthening of the infectious period [35]. Fever, although invalidating for the patient, reduces pathogen replication rates and enhances the adaptive immune response [36]. It has even been suggested that the use of aspirin during the 1918 outbreak might have caused an increase in mortality [37]. Even in the case of non-lethal infection, we can hypothesize that TOL, by increasing contact rates, will raise transmission and thus the incidence and prevalence of the infection.

Tolerance treatments might, however, be applied in hospitals, where transmission can be curbed. Of particular interest are antivirulence drugs [38, 39], because of their potential for the management of antimicrobial resistance. Tolerance mechanisms involving free heme regulation also received broad interest in the last years, as they suggest promising applications for treating severe sepsis [13], especially since available treatments are limited [40]. The targeting of free circulating heme, which contributes to pathogenesis indepently of pathogen load, is a case of tolerance-based treatment that can be applied after bacteria have been cleared. The treatment we discussed above may be applicable not only because transmission can be controlled in a hospital setting, but also because sepsis is treated after bacterial clearance.

Our results suggests that pathogen evolution could have a dramatic effect in chronic infections, where infected hosts are infectious for a long time. The mortality, that already increases because of the epidemiological feedback, will be amplified by the higher virulence of evolved pathogens. Further studies are needed to assess the public health implications of pathogen evolution in response to TOL. To go beyond the insights of Vale et al., such studies should focus on the non-prophylactic use of TOL and consider a broad range of treatment coverage.

The main disadvantage of TOL that we identified does not rely on the evolution of the pathogen but arises simply through the epidemiological feedback that is amplified by TOL. Our results raise serious doubts about the promise of tolerance-based strategies, especially when treating chronic infections.

## Supporting Information

- **S1 text.** Equilibrium analysis of the tolerance-based treatment model.
- **S2 text.** Equilibrium analysis with pathogen resistance to treatment.
- **S3 text.** Population-level impact of imperfect tolerance-based treatment.
- **S4 text.** Why are the population-level effects more pronounced for chronic than for acute infections?
- **S5 text.** Comparison of prophylactic and post-exposure treatments.
- **S6 text.** Impact of the different treatments on the basic reproduction number.
- **S7 text.** Can we curb disease incidence with transmission control?
- **Fig S1.** Robustness of the results to parameter changes.
- **Fig S2.** Effect of *R*_0_ on the severity of infection.
- **Fig S3.** Effect of the relative values of *v*_*TOL*_ and *v* on the disease-induced mortality.
- **Fig S4.** Effect of transmission control on prevalence, incidence and mortality.

